# Complex multi-trait responses to multivariate environmental cues in a seasonal butterfly

**DOI:** 10.1101/772749

**Authors:** Pragya Singh, Erik van Bergen, Oskar Brattström, Dave Osbaldeston, Paul M. Brakefield, Vicencio Oostra

## Abstract

Developmental plasticity in a seasonal environment allows an organism to optimally match its life-history traits with the fluctuating conditions. This critically relies on abiotic and biotic factors, such as temperature or food quality, that act as environmental cues and predict seasonal transitions. In most seasonal environments, multiple factors vary together, making it crucial to understand their combined effects on an organism’s phenotype. Here, we study plasticity in a multivariate environment in the butterfly *Bicyclus anynana* that exhibits two distinct seasonal phenotypes. Temperature is an important cue mediating plasticity in this species, but other environmental cues such as larval host plant quality could also be informative since plant quality deteriorates during the transition from wet to dry season in the field. We examine how temperature and host plant quality interact to affect life-history traits. Using a full-factorial design, we expose cohorts of larvae to either poor (old plants) or high (young plants) quality plants at different temperatures. Our results show that plant quality had a temperature and sex-dependent effect on life-history traits. At lower and intermediate temperatures, it decreased body mass and prolonged development time, indicating that poor plant quality acted as a stressor. However, metabolic rates in adults were not affected, indicating that individuals could, at least in part, compensate for stressful juvenile conditions. In contrast, at higher temperatures poor plant quality induced a partial dry-season phenotype, indicating that it may have acted as an environmental cue. Moreover, poor plant quality, particularly in males, also decreased the correlation between life history traits, signifying disrupted phenotypic integration. Our study reveals complex interactive effects of two environmental variables on seasonal plasticity, reflecting differences in their reliability as seasonal cues. This highlights the importance of studying the combined effects of multiple environmental factors to better understand the regulation of phenotypic plasticity in wild.

## Introduction

Environmental seasonality is frequent in nature and can lead to the evolution of phenotypic plasticity (Tauber et al. 1986, Gotthard and Nylin 1995, Lafuente and Beldade 2019). Plasticity can help to ensure that the phenotype expressed by an organism matches the requirements of the organism’s environment, and often involves trade-offs between life-history traits such as reproduction and lifespan, or growth rate and body size (Nylin 1992, Flatt and Heyland 2011, Torres-Dowdall et al. 2012). A particular case of phenotypic plasticity is developmental plasticity, where phenotypic changes are induced by the environment experienced during development (Beldade et al. 2011). Developmental plasticity can be adaptive in seasonal environments as it can allow organisms to adjust their life history strategy for future conditions well before the new season starts, using predictive environmental cues present during the course of development. Environmental cues can range from abiotic factors, like temperature and day-length, to biotic factors, such as food quality and predator presence. A special type of developmental plasticity is the Predictive Adaptive Response (PAR), in which an organism adjusts its phenotype to maximize fitness in future environmental conditions, even though the phenotype might not be immediately advantageous in the current environment. In fact, the phenotype could be maladaptive if the expected environmental change does not occur (Ghalambor et al. 2007, Monaghan 2008, Saastamoinen et al. 2010).

In a seasonal environment multiple environmental factors often vary together (Jackson et al. 2009, Chevin and Lande 2015). For example, low precipitation is usually accompanied by a decline in quantity or quality of resources. This leads to key open questions about i) whether organisms sense the environment through one or multiple cues, ii) whether these cues interact (E × E) or act as independent predictors, and iii) whether they are processed in similar manners leading to similar phenotypic responses. One hypothesis would be that the factors act together as cues to predict future environmental conditions and potentially lead to an additive adaptive response. Alternatively, one factor could act as a cue, while the other environmental factor may not be perceived or processed at all, or could even be a stressor that disrupts the phenotype or phenotypic integration. Theoretical work has shown that when the relationship between an environmental cue and future conditions is weak, plasticity may not evolve in response to the environmental predictor (Tufto 2000, Leimar et al. 2006, Reed et al. 2010). Moreover, in a case where multiple cues are used by an organism to respond to the environment, the responses to a single cue might be nonintuitive and misleading (Chevin and Lande 2015). While many earlier studies on plasticity have manipulated one environmental cue at a time, it is more likely that multiple environmental cues vary simultaneously in natural seasonal environments, making it important to study plasticity in a multivariate environment.

To investigate the effect of multivariate environment on developmental plasticity, we use the seasonally polyphenic butterfly *Bicyclus anynana*. This species exhibits two alternative seasonal forms (dry and wet) which correspond to a cool and a warm season, respectively. In addition to differences in morphology (Brakefield and Reitsma 1991), physiology (Oostra et al. 2011), and behaviour (van Bergen and Beldade 2019), the forms also differ markedly in their life-history traits (e.g. Oostra et al. 2011). Previous studies have shown that the temperature experienced during the (late) larval stage is a crucial cue for plasticity in this species (Brakefield and Reitsma 1991, Brakefield et al. 2007, 2009). Interestingly, variation in temperature alone does not produce the full extent of plasticity in life-history traits as observed in the wild (Roskam and Brakefield 1999), suggesting that other predictive environmental factors may act in conjunction with temperature (Brakefield 1987, Brakefield and Reitsma 1991). Here, we hypothesise that larval host plant quality could be an important environmental cue, in addition to temperature, for developing individuals in the field. During the transition from wet to dry season in the field, the plants on which the larvae feed tend to be old and of poor quality (Brakefield and Reitsma 1991, Kooi et al. 1996). Moreover, earlier work in *B. anynana* has shown that this species, under conditions of food limitation, is able to produce a PAR phenotype (i.e. better adapted to cope with stressful conditions as an adult) (Saastamoinen et al. 2010, van den Heuvel et al. 2013).

In our study, we test how larval host plant quality–in conjunction with temperature–affects a suite of life history traits. In particular, using old host plants that mimic the deteriorating conditions in dry season, we feed cohorts of individuals for a part of larval development on old (poor quality) plants. Data collected from these individuals is compared to those of cohorts that had been reared on young (high quality) plants throughout development. These treatments enable an examination of whether larval host plant quality affects the phenotype. Furthermore, we test the effect of host plant quality at three different temperatures that correspond to wet, intermediate and dry season temperatures in the field. This design allows testing of how larval host plant quality and temperature interact to affect larval and pupal development time, pupal and adult mass, resting metabolic rate (RMR) and the respiratory quotient of adults. Earlier studies using this species examined RMR by measuring CO_2_ respiration rate (Brakefield et al. 2007, Pijpe et al. 2007), and have showed that it varies in response to temperature. However, as of yet, no study has examined the O_2_ consumption or respiratory quotient in *B. anynana*. Examining the respiratory quotient allows us to evaluate whether adults differ in their macronutrient metabolism in response to environmental conditions (i.e. whether they burn different fuels, in particular fat, protein and carbohydrates). Finally, we tested whether the host plant quality affects the organismal integration of phenotypic traits by examining the correlation between life-history traits. The response of different phenotypic traits to environmental cues might be correlated (van Bergen et al. 2017), e.g. due to similar underlying physiological processes, but under stress we might expect the synchronised response of the traits to become uncoupled and phenotypic integration to decrease.

## Materials and Methods

### Study organism

*Bicyclus anynana* is a Nymphalid butterfly from East Africa and is a model organism for studying seasonal and developmental plasticity (Brakefield et al. 2009). It is found in savannah grasslands and open woodlands (both seasonal ecosystems) and has probably evolved developmental plasticity as an adaptation to seasonality in the environment. The two seasons that *B. anynana* experiences are the warm wet season and the cool dry season, and the species expresses alternative morphs in these two alternative seasons. The wet season butterflies experience high temperatures and precipitation during development, and adults have larger, more conspicuous eyespots on their ventral wing surfaces, shorter larval and pupal developmental periods, lower pupal and adult mass, shorter lifespan and reproduce relatively early (Brakefield et al. 2009, Oostra et al. 2011). In contrast, the dry season larvae that develop during the late wet season experience relatively cooler temperatures and lower precipitation which is associated with the transition from wet to dry season (Windig et al. 1994, van Bergen et al. 2016). Moreover, dry season adults accumulate higher mass and fat reserves during development; have small or absent eyespots, a higher metabolic rate, delayed reproduction until the following wet season, and a longer lifespan (Brakefield and Reitsma 1991, Pijpe et al. 2007, Oostra et al. 2011, Halali et al. 2019). The seasons, along with differing in temperature and precipitation, also differ drastically in the availability of resources, with the cool dry season having a reduced host plant quantity and quality (Roskam and Brakefield 1999, van Bergen et al. 2016). The adults of this butterfly species feed on rotting and fermenting fruit and the larvae utilize grasses.

### Experimental design and rearing

An outbred laboratory stock of the butterfly *B. anynana* was used for the experiment. The stock was established in 1988 from numerous gravid females collected in Malawi. Adults are fed on banana, and the larvae are reared on maize (*Zea mays*) (Brakefield et al. 2009).

We used a full-factorial design to investigate the effects of larval host plant quality, pre-adult (i.e. larval and pupal) temperature, and their interaction on a suite of life-history traits. Three temperature treatments (19, 23 and 27°C, representing dry, intermediate, and wet season conditions, respectively) and two plant quality treatments (old maize, young maize) were used. Eggs were collected daily from the stock population. One day after hatching, larvae were randomly allocated to cages (35cm × 44cm × 65cm) with young maize plants set up in climate rooms at 19°C and 27°C, and in smaller climate-cabinets (Sanyo/Panasonic MLR-350H) at 23°C (all at 75% relative humidity and a 12h:12h day: night light cycle). Each cage had at least 140 larvae (950 larvae in total, Table 1). One day after they moulted to the 4th instar, larvae were switched to the host plant treatment: old maize or fresh young plants. The larvae were only exposed to the host plant treatment during the final two larval instars, which is the period when most growth occurs and the effect of food quality should be most prominent. In addition, this period is close to the pupal stage when the adult phenotype is differentiated. The resulting pupae were then individually placed in transparent pots, assigned an ID and kept at their temperature treatment, until they eclosed.

**Table 1.**
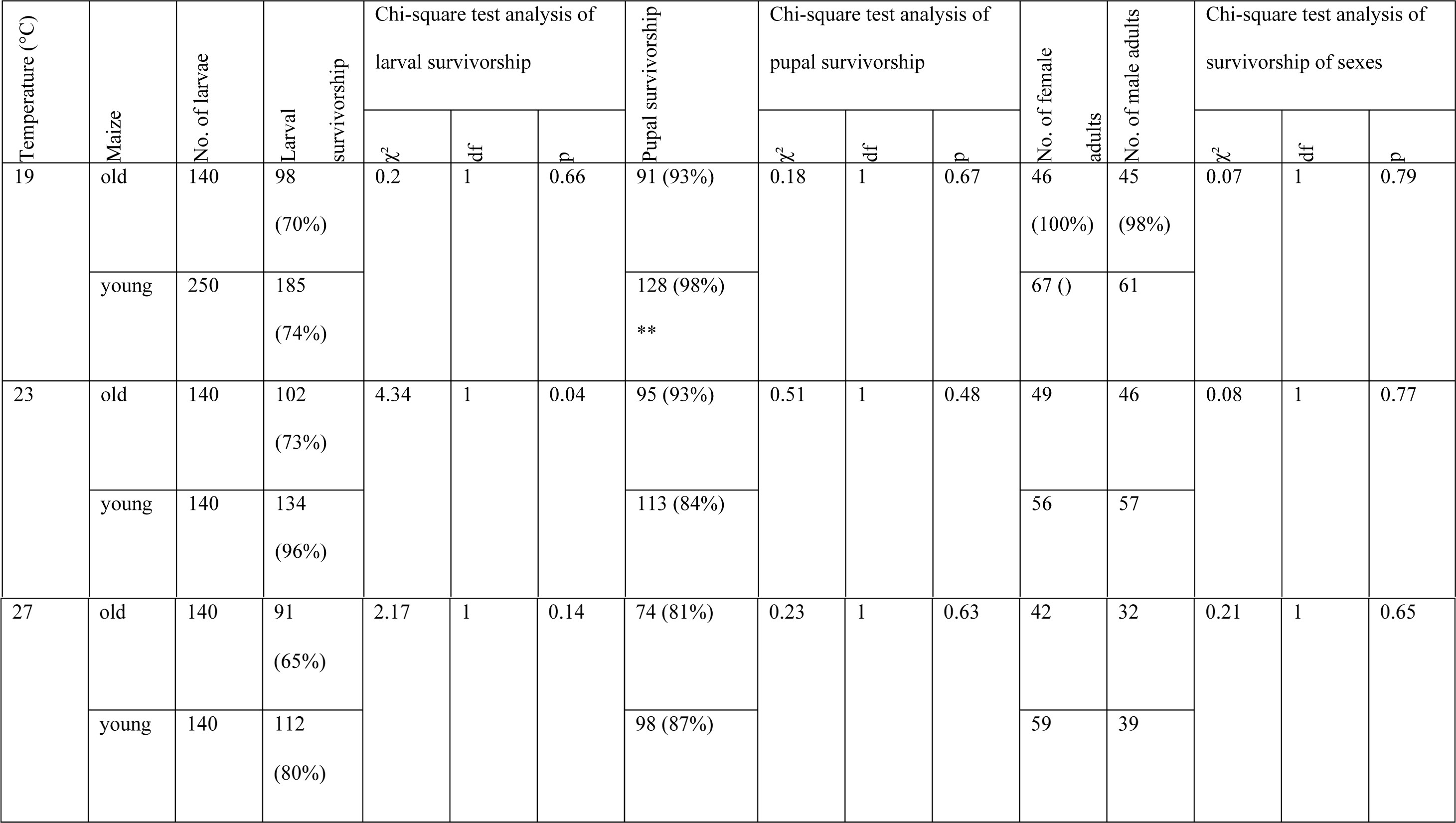
Larval and pupal survival (% in brackets) for different temperatures. The larval survivorship was computed as the number of pupations and is expressed as a percentage of the initial number of larvae reared. Pupal survivorship was the number of adults which eclosed and is expressed as a percentage of the initial number of pupae. * * We discarded excess pupae at 19°C and only kept 130 pupae.

| Temperature (°C) | Maize | No. of larvae | Larval survivorship | Chi-square test analysis of larval survivorship |  |  | Pupal survivorship | Chi-square test analysis of pupal survivorship |  |  | No. of female adults | No. of male adults | Chi-square test analysis of survivorship of sexes |  |  |
| --- | --- | --- | --- | --- | --- | --- | --- | --- | --- | --- | --- | --- | --- | --- | --- |
| | | | | $\chi^2$ | df | p | | $\chi^2$ | df | p | | | $\chi^2$ | df | p |
| 19 | old | 140 | 98<br>(70%) | 0.2 | 1 | 0.66 | 91 (93%) | 0.18 | 1 | 0.67 | 46<br>(100%) | 45<br>(98%) | 0.07 | 1 | 0.79 |
|  | young | 250 | 185<br>(74%) |  |  |  | 128 (98%)<br>** |  |  |  | 67 () | 61 |  |  |  |
| 23 | old | 140 | 102<br>(73%) | 4.34 | 1 | 0.04 | 95 (93%) | 0.51 | 1 | 0.48 | 49 | 46 | 0.08 | 1 | 0.77 |
|  | young | 140 | 134<br>(96%) |  |  |  | 113 (84%) |  |  |  | 56 | 57 |  |  |  |
| 27 | old | 140 | 91<br>(65%) | 2.17 | 1 | 0.14 | 74 (81%) | 0.23 | 1 | 0.63 | 42 | 32 | 0.21 | 1 | 0.65 |
|  | young | 140 | 112<br>(80%) |  |  |  | 98 (87%) |  |  |  | 59 | 39 |  |  |  |

After randomly culling excess pupae raised at 19°C and excluding 51 individuals due to missing information about one or multiple life-history traits, the final sample size across all treatments was 548 (see Table 1).

### Host plant quality treatments

All maize plants were grown from seed and reared in a climate-controlled greenhouse in Madingley (United Kingdom), with regular watering to keep the soil moist at all times. Young maize plants were 2-3 weeks old whereas old maize plants were at least 5-7 weeks old. Earlier studies have shown that plant quality varies with age. For example, older plants typically have tougher leaves (Choong 1996, Loney et al. 2006), lower nutritional values (Hikosaka et al. 1994) and different chemical/physical defences against herbivory (Barton and Koricheva 2010) than younger plants. These differences can have pronounced effects on herbivory (Price et al. 1987, Loney et al. 2006), with the incidence of herbivorous invertebrates on old host plants typically being lower than on young plants (Choong 1996, Fenner et al. 1999, Boege and Marquis 2005). Thus, older host plants can be inferred to be of poor quality relative to younger host plants. Moreover, host plant quality can also directly regulate phenotypic plasticity in herbivorous insects (Lin et al. 2018).

In our experiment, we measured the maximum leaf width and height of each maize plant before feeding it to the larvae. For old maize plants, plant height was 92.2±33.2 (mean±sd) cm and maximum leaf-width was 4.2±0.6 cm. For young maize plants, plant height was 69.6±4.5 cm and maximum leaf-width was 1.4±0.2 cm. The larvae were reared on whole plants (multiple larvae per plant), and *ad libitum* feeding was ensured by providing new plants whenever needed. When the old plants were too large to be completely accommodated inside the cage, only a part of the (whole) plant was put inside, while ensuring that the larvae could not escape from the cage.

### Life-history traits

For each individual, larval development time was recorded as the number of days between hatching of the egg and pupation of the larvae, and pupal development time was recorded as the number of days between pupation and eclosion of the butterfly. Pupae were weighed approximately 24 h after pupation. Adults were weighed and resting metabolic rate (RMR) measurements made one day after eclosion. For the RMR measurements, individual butterflies were measured in the dark–at their rearing temperature–in small cylindrical glass containers (4 cm in diameter × 9 cm in height). Each RMR cycle consisted of three runs of 20 minutes during which RMR was measured as the individual rate of CO_2_ and O_2_ respiration (millilitre per minute), using stop-flow respirometry (Pijpe et al. 2007). CO_2_ and O_2_ production were measured using a LI-7000 CO_2_ gas analyser (Li-Cor) and an Oxzilla FC-2 Differential Oxygen Analyzer (Sable Systems), respectively, and acquired data were handled in Expedata (Sable Systems). The CO_2_ and O_2_ respiration rates were scaled to mass by dividing respiration rate by adult mass. Measurements were taken at the same time of the day for all individuals, and the data from the second and third runs were averaged. The first run was excluded for each individual as this occurred during the butterfly’s acclimation phase. The respiratory quotient was calculated as the CO_2_ respiration rate divided by the O_2_ respiration rate (Richardson 1929).

### Statistical analyses

We performed a Chi-square Goodness-of-Fit test to test if host plant quality had an effect on the larval and pupal survivorship. We also performed a Chi-Square Test for Independence to assess if the host plant quality had a sex-specific effect on survivorship for each temperature. In addition, for each dependent variable (larval development time, pupal development time, pupal mass, adult mass, CO_2_ and O_2_ respiration rates (unscaled and scaled by mass), and respiratory quotient), we constructed a three-way ANOVA with temperature, host plant quality, sex, and all interactions, as independent fixed factors. We evaluated each model and removed the least significant term from this model at 0.05 significance level (p<0.05) in a step-wise fashion to obtain the minimal adequate model. Here we report the results of the minimal adequate model for all the dependent variables (Supplementary S1). Prior to statistical analyses, all traits, except pupal mass, were log-transformed when they did not fulfil the assumptions of a parametric test. We did post-hoc comparisons using t-tests and corrected for multiple testing with Bonferroni correction.

To assess whether host plant quality had an effect on phenotypic integration, we calculated Pearson’s correlation coefficients among the log-transformed life-history traits (all traits except unscaled CO_2_ and unscaled O_2_ respiration rates) for individuals reared on both young and old host plants, and for each sex (i.e. four correlation coefficients per pair of traits). We tested whether poor host plant quality disrupted the seasonal morphs by comparing the correlation coefficient of each life-history trait pair between old and young host plants. We converted the correlation coefficient into a *z*-score using Fisher’s *r*-to-*z* transformation and compared these *z*-scores using the sample size for each coefficient, using the following formula (Cohen et al. 2003):

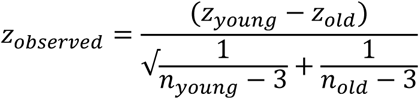

where z_young_ and z_old_ are correlation coefficients and n_young_ and n_old_ are the sample sizes for individuals on young and old host plants, respectively. We assessed whether the difference between the correlation coefficients was statistically significant (P ≤0.05) by checking if the z_observed_ was greater than the critical value of ± 1.96.

All the analyses were done in R version 3.1.0 (R Development Core Team 2016).

## Results

In general, temperature and sex had a significant effect on all traits except respiratory quotient, as had been found in previous studies (Pijpe et al. 2007, de Jong et al. 2010, Oostra et al. 2011, 2014, Mateus et al. 2014). The effect of host plant quality varied among traits, typically with a temperature and/or sex dependent effect (Table 1, Supplementary S1).

### Limited effect of host plant quality on pre-adult survivorship

Development on old maize had a limited effect on larval and pupal survivorship, with a significant effect only on larval survivorship at 23°C, where survival was higher on young maize plants (Table 1). Moreover, host plant quality did not have a sex-specific effect on survivorship (Table 1).

### Prolonged development at 23°C due to poor host plant quality

Consistent with earlier studies (Pijpe et al. 2007, de Jong et al. 2010, Oostra et al. 2011, Mateus et al. 2014), development time decreased with increase in temperature, and males had a shorter larval but longer pupal development time than females (Supplementary S1). In contrast to the treatments at both ends of the thermal gradient (19°C and 27°C), host plant quality had a significant effect on larval and pupal development time at 23°C (Figure 1). At this intermediate temperature the larvae/pupae took a longer time to develop on old maize.

**Figure 1.**
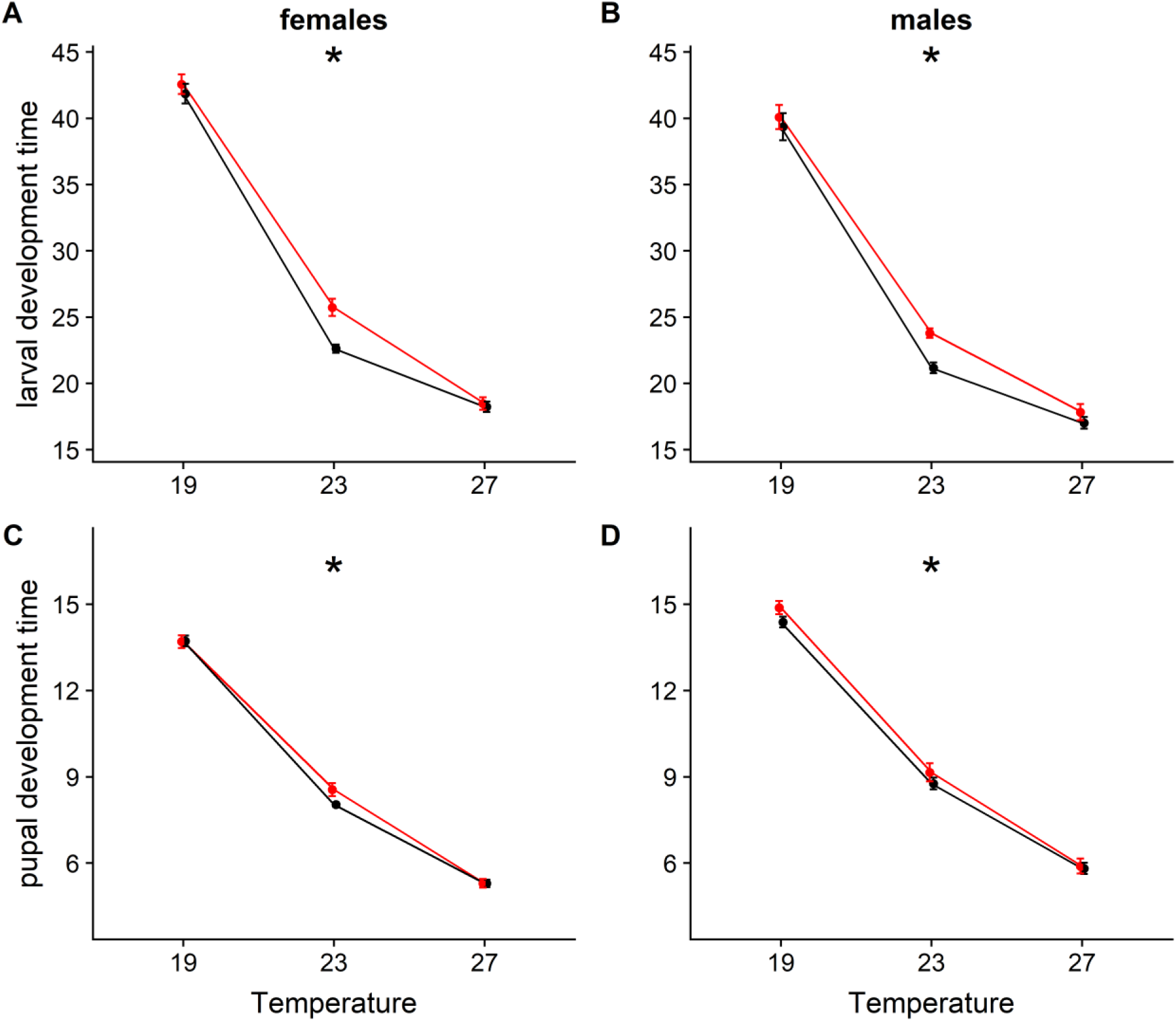
Slower development due to poor host plant quality at certain temperatures: Effect of host plant quality and temperature on larval development time (top row) and pupal development time (bottom row) in each sex. Plots show means and 95% confidence intervals of data. Statistically significant effect of host plant quality on development time is indicated for each temperature with *, indicating P≤0.05. Red lines represent old host plants and black lines represents young host plants.

### Temperature-dependent effects of host plant quality on body mass

Host plant quality led to a temperature-specific reduction in pupal mass, and to a temperature and sex-specific effect on adult mass (Supplementary S1, Figure 2). On old maize, both sexes had lower pupal mass at 19°C and 23°C, while both sexes had higher pupal mass on old maize at 27°C, though the latter was not statistically significant. Furthermore, pupal mass significantly decreased with increasing temperature, and was higher in females. Both sexes had lower adult mass on old maize at 19°C, while only females had a lower adult mass on old maize at 23°C. Similar to pupal mass, both sexes had higher adult mass on old maize at 27°C, though this effect was only significant for females. Females had a higher adult mass compared to males for all three temperatures.

**Figure 2.**
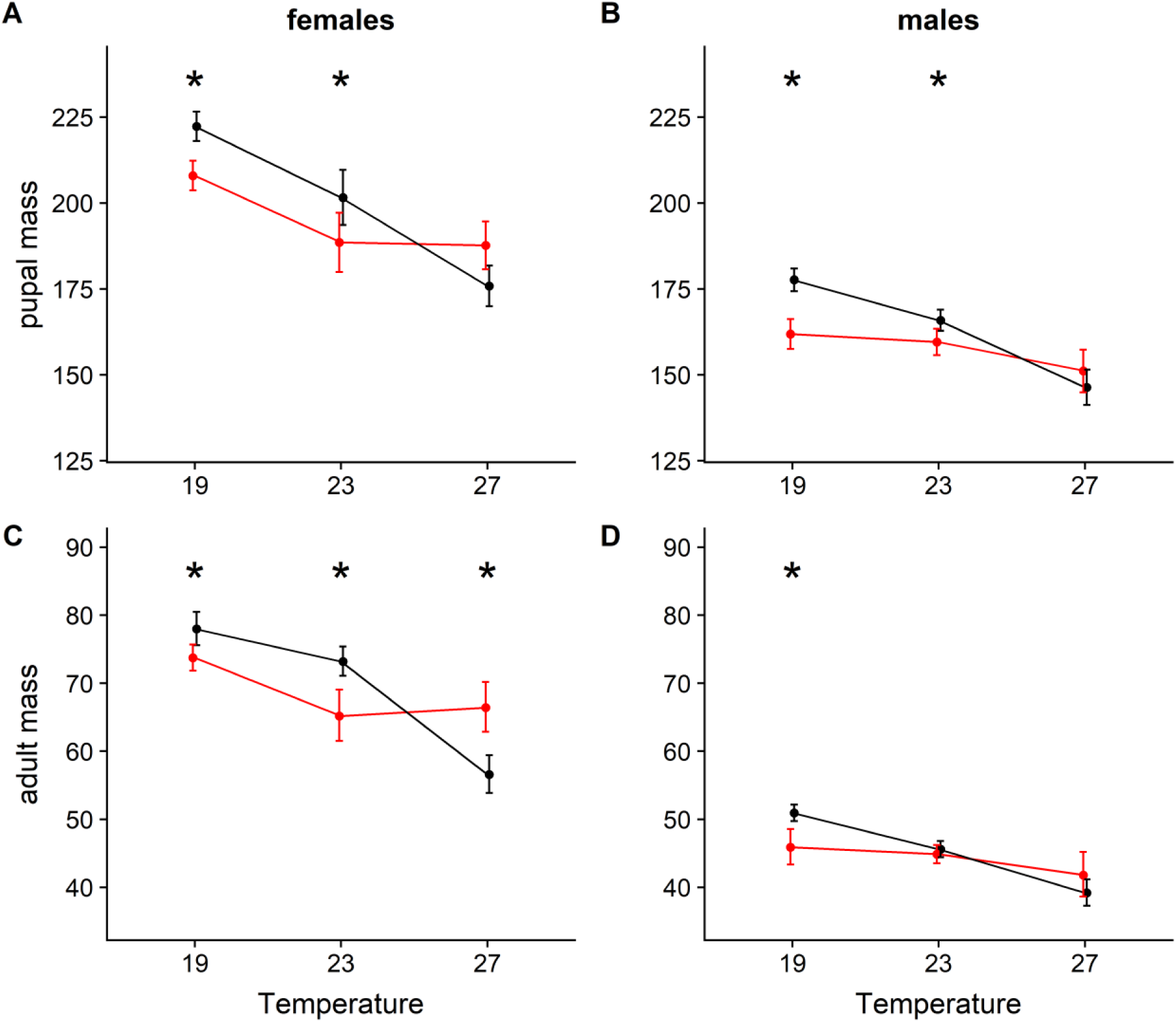
Temperature and sex-dependent effects of host plant quality on body mass: Effect of host plant quality and temperature on pupal mass (top row) and adult mass (bottom row) in each sex. Plots show means and 95% confidence intervals of data. Statistically significant effect of host plant quality on development time is indicated for each temperature with *, indicating P≤0.05. Red lines represent old host plants and black lines represents young host plants.

### No effect of host plant quality on mass-scaled respiration rates and respiratory quotient

Similar to pupal mass, unscaled CO_2_ and O_2_ respiration rate were significantly affected by the interaction between temperature and host plant quality (Supplementary S1, Figure 3, 4). Adults of both sexes had a higher unscaled CO_2_ respiration rate when reared on old compared to young maize at 27°C (Figure 3). In contrast, there was no significant effect of plant quality on unscaled O_2_ respiration rate at any temperature (Figure 4). Females had significantly higher unscaled CO_2_ and O_2_ respiration rates than males. Moreover, on both old and young plants, unscaled CO_2_ and O_2_ respiration rates were significantly lower at 19°C compared to the other temperatures, but did not differ significantly between 23°C and 27°C. However, after scaling (correcting) for body mass, only temperature and sex had a significant effect on CO_2_ and O_2_ respiration rates (Supplementary S1, Figure 3, 4), indicating that the effect of food stress on CO_2_ and O_2_ respiration was modulated by a difference in body mass. Similar to earlier studies on CO_2_ respiration rates in this species (Brakefield et al. 2007, Pijpe et al. 2007), both the CO_2_ and O_2_ respiration rate increased with temperature (with all temperatures being significantly different from each other), and males had higher respiration rates than females. The respiratory quotient was not affected by sex, temperature, food stress, or any interactions between these factors (Supplementary S1, Figure 5).

**Figure 3.**
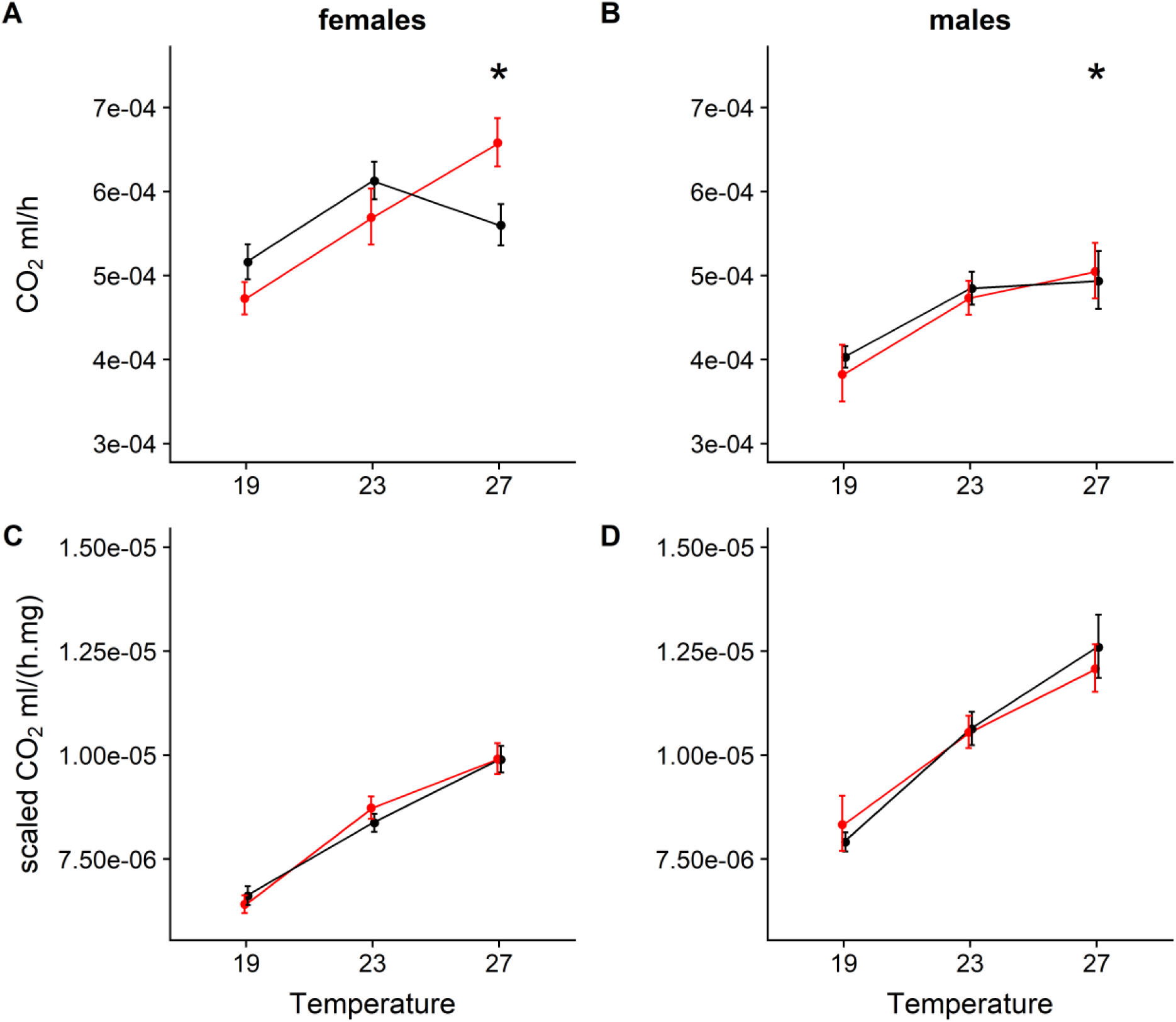
Limited effect of host plant quality on unscaled and scaled CO_2_ respiration rates (production per hour): Effect of host plant quality and temperature on unscaled CO_2_ respiration rate (top row) and CO_2_ respiration rate scaled by mass (bottom row) in each sex. Plots show means and 95% confidence intervals of data. Statistically significant effect of host plant quality on development time is indicated for each temperature with *, indicating P≤0.05. Red lines represent old host plants and black lines represents young host plants.

**Figure 4.**
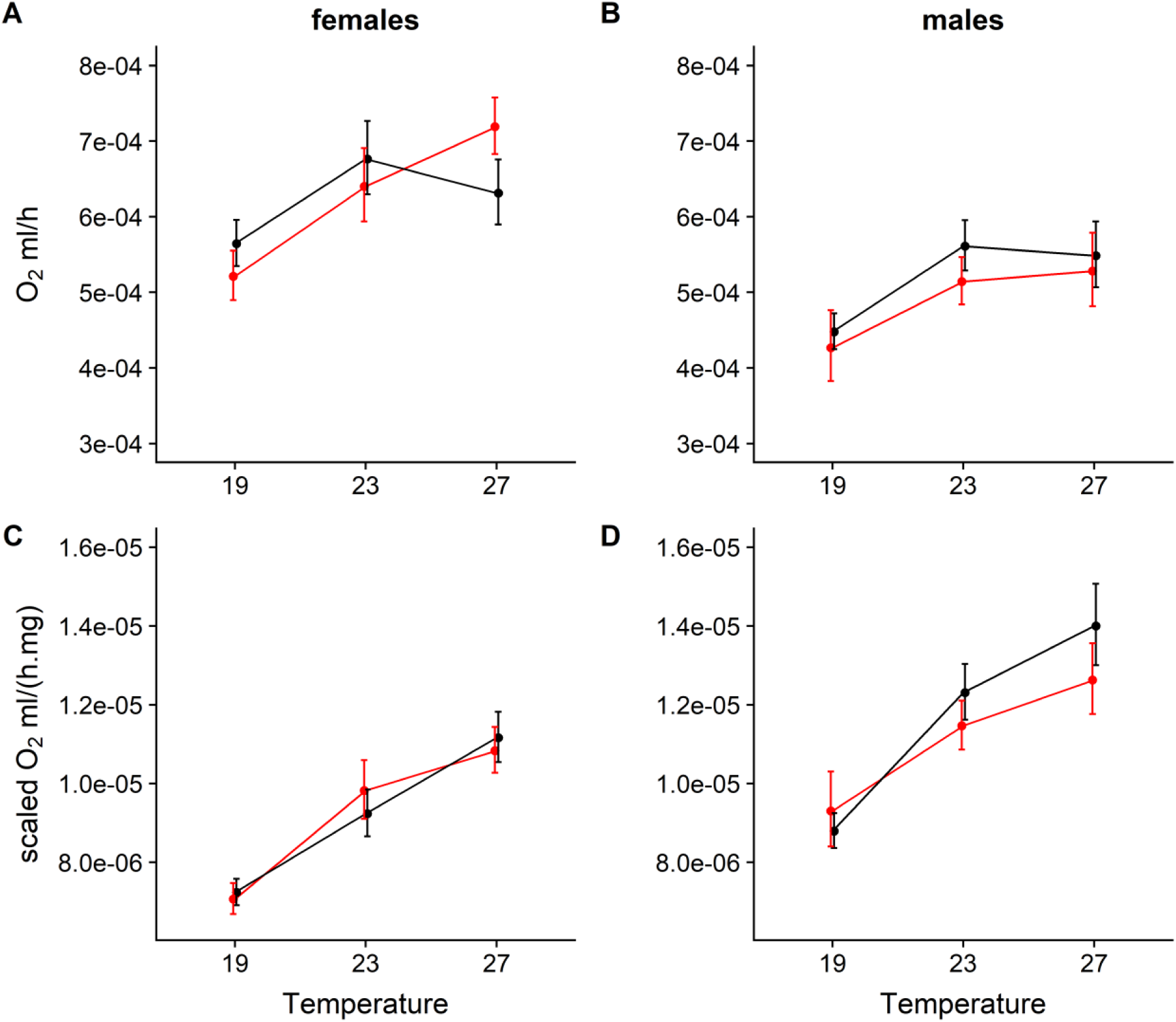
No effect of host plant quality on unscaled and scaled O_2_ respiration rates (production per hour): Effect of host plant quality and temperature on unscaled O_2_ respiration rate (top row) and O2 respiration rate scaled by mass (bottom row) in each sex. Plots show means and 95% confidence intervals of data. Red lines represent old host plants and black lines represents young host plants.

**Figure 5.**
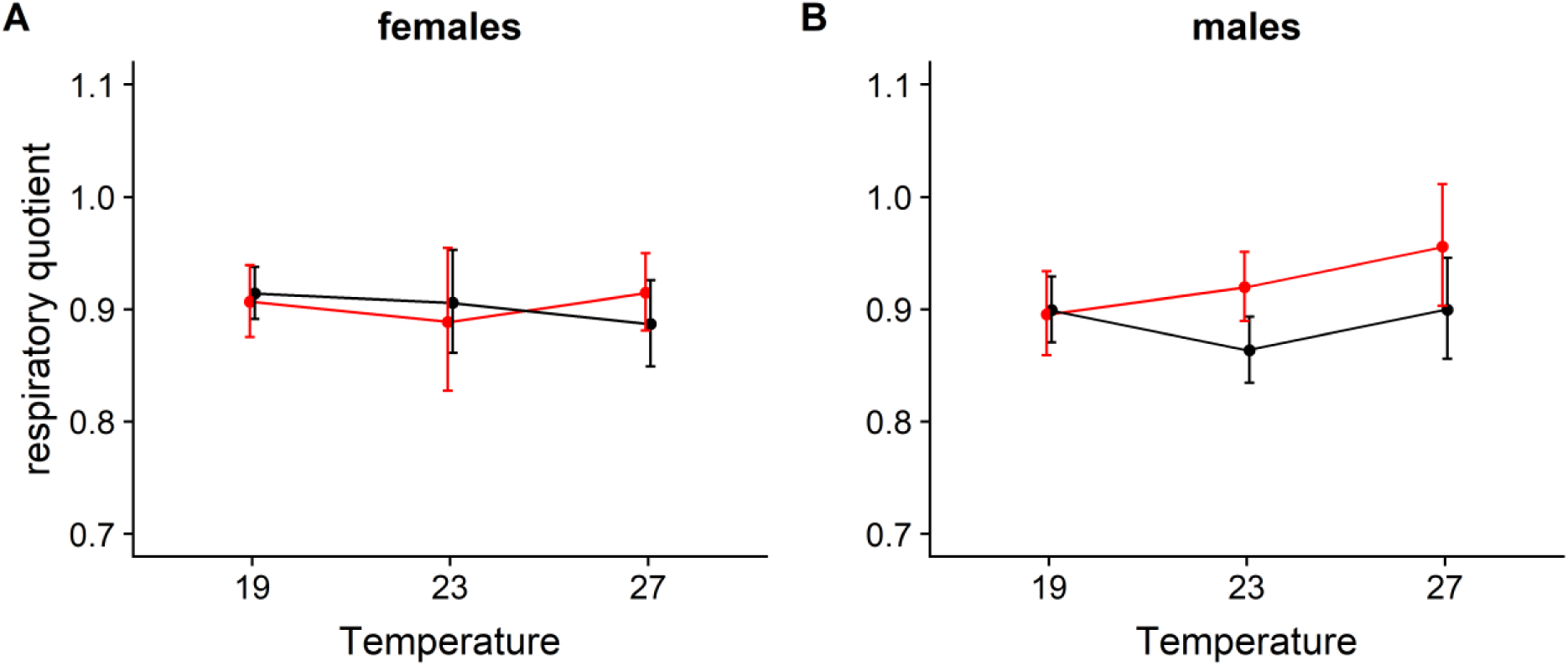
No effect of temperature and host plant quality on respiratory quotient in each sex. Plots show means and 95% confidence intervals of data. Red lines represent old host plants and black lines represents young host plants.

### Poor host plant quality disrupts phenotypic integration

Host plant quality significantly changed the correlation matrix for life-history traits in both sexes (Figure 6, for details see Supplementary S2). Males were more severely affected, with 13 out of 21 correlation coefficients being significantly different between young and old host plants, while for females only 7 out of 21 correlation coefficients were significantly affected. Amongst the significant changes, for males, all 13 correlation coefficients decreased on old host plants while for females 5 correlation coefficients decreased and 2 correlation coefficients increased on old host plants. Thus, in general, poor host plant quality resulted in a less integrated phenotype with a decrease in correlations among life-history traits, especially for males.

**Figure 6.**
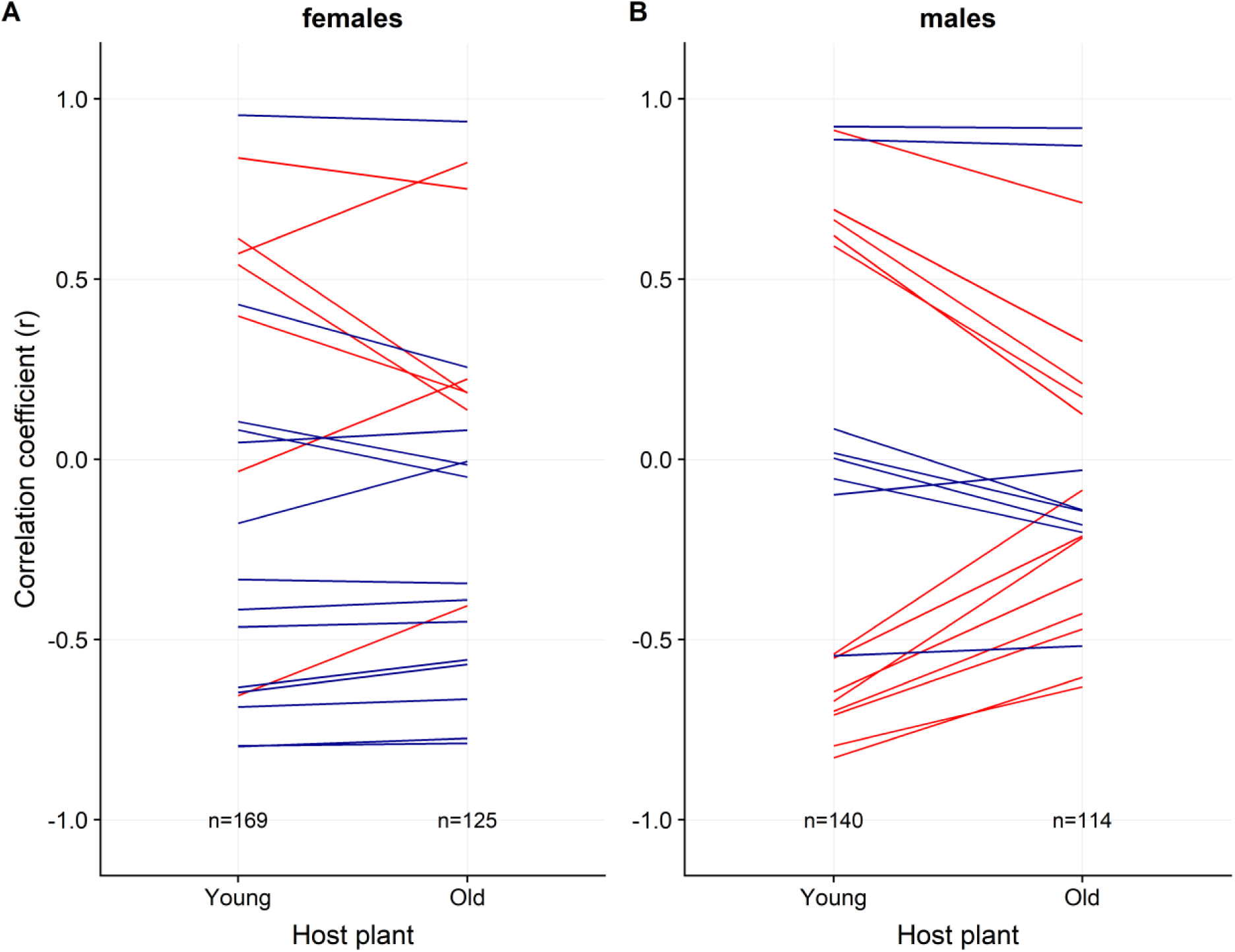
Poor host plant quality disrupts phenotypic integration, particularly for males: Correlation coefficients (r) between log transformed trait values for a) females and, b) males on old (poor quality) or young (high quality) host plants. Each line represents the correlation coefficient between one pair of traits. Correlation coefficients that changed significantly (P≤0.05), due to poor host plant quality, are highlighted in red. Sample sizes for calculating each correlation coefficient are given at the bottom.

## Discussion

While several earlier studies have examined the effect of food stress on life-history traits in different organisms, relatively few studies have analysed the role of food quality as a predictive cue for future conditions, especially in combination with other cues such as temperature (Wilson 1994, Rotem et al. 2003, Bauerfeind and Fischer 2005, Coley et al. 2006, Rosa and Saastamoinen 2017). Examining the effect of food quality in combination with other environmental cues is important, as most seasonal environments in nature are usually characterized by multiple environmental factors varying together. In our experiment, we examined the effect of host plant quality in conjunction with temperature, and found that host plant quality had a temperature- and sex-specific effect on life-history traits.

Earlier studies using *B. anynana* and food deprivation as nutritional stress, had shown that stressed individuals had a reduced body mass and prolonged developmental time (Bauerfeind and Fischer 2005, Saastamoinen et al. 2010, 2013). These results contrast our findings and the difference in effect is likely due to the different approaches used; in the earlier studies individuals were starved (food deprivation) while in our study individuals were fed *ad libitum* on old host plants. These older host plants are expected to be less palatable (i.e. tougher), have lower nutritional values and differ in their composition of secondary metabolites relative to younger plants. Compared to food deprivation, larvae developing on old host plants were likely able to acquire sufficient resources and thus, demonstrated a less dramatic change in the life-history traits studied. Moreover, food deprivation and old host plants may activate different physiological responses.

In our study, the effect of host plant quality on different life-history traits was temperature-dependent, indicating that the effect depended on the physiological state of the organism (which can also depend on temperature experienced). If reduced host plant quality can be perceived as an environmental cue indicative of harsh (future) environmental conditions, we would have expected development on old host plants to lead to a more dry-season phenotype (e.g. an increase in body mass). Indeed, when exposed to the thermal conditions of the wet-season (27°C), poor host plant quality induced a partial dry-season phenotype with an increase in adult mass (significant for females). In contrast to the pattern observed at high temperatures, at temperatures that mimic the dry season (19°C), poor host plant quality did not act as a seasonal cue. Instead, the treatment resulted in lower body mass and longer development times, indicating a stress response. The increase in adult mass in wet-season form butterflies (27°C) could potentially indicate an adaptive response (e.g. PAR-like) for within-season fluctuations in food quality. Food quality can potentially vary independently of temperature, making it an important cue under conditions when the thermal information may be inconclusive (see below).

Interestingly, for several traits, including larval survivorship and development time, we observed a significant effect of host plant quality only at 23°C, which is the average temperature during the transition from the wet (27°C) to the dry (19°C) season (Windig et al. 1994, van Bergen et al. 2016). One possible explanation for observing the above effects only at 23°C could be that increased sensitivity for host plant quality could be adaptive at this temperature. Larvae developing during the transition between the seasons, or during a cold spell in the wet season, may require additional environmental information to induce the expression of the appropriate phenotype (see above). Moreover, the prolonged development time at 23°C is likely due to the old maize being of a poorer quality (e.g. tougher leaves and/or more secondary metabolites), prolonging the period necessary to reach the critical mass needed for undergoing hormonal changes and pupation (Coley et al. 2006). Remarkably, despite the prolonged time to acquire resources, the individuals reared on old plants had a lower pupal mass (both sexes) and adult mass (only significant for females).

In addition, the effect of host plant quality on body mass was more evident compared to the effect on development time and survivorship at all thermal environments. This may be related to the fact that the larvae were only exposed to the poor host plant quality during the final two larval instars. The latter represents the period when most growth occurs, but it is only a short period of the total development time, potentially resulting in more pronounced effects on body mass compared to development time.

We also examined for the first time the respiratory quotient, the ratio between CO_2_ and O_2_ respiration rate at rest. We found that it was not influenced by either temperature, sex, host plant quality, or their interactions. The respiratory quotient reflects which macronutrients are metabolized for energy, with values of 0.7, 0.8 or 1.0 indicating fat, protein or carbohydrate metabolism, respectively (Nunes et al. 1997). In our study, the respiratory quotient was around 0.9, which is intermediate between protein and carbohydrate metabolism, across all temperatures. Thus, our results point towards adult macronutrient metabolism remaining similar, irrespective of larval food quality. This is surprising, as earlier studies in both field and laboratory showed that dry season form butterflies have a higher fat content (Brakefield and Reitsma 1991, Pijpe et al. 2007, de Jong et al. 2010, Oostra et al. 2011). However, we measured the metabolic rates of newly eclosed adults under benign conditions in the laboratory where fat reserves are likely under-used compared to the wild, where adults often face prolonged periods of desiccation and/or starvation.

The overall phenotypic integration of traits was affected by plant quality, with a decrease in the correlation between life history traits on poor quality host plants, signifying disrupted phenotypic integration, particularly in males. Interestingly, a similar phenomenon of sex-specific effects of larval food stress on larval and adult performance has been found in a study using the Glanville fritillary butterfly, *Melitaea cinxia* (Rosa and Saastamoinen 2017). This could indicate that the underlying physiological processes that are responsible for phenotypic integration, e.g. hormone signalling, may have a sex-specific component, and that stressors affecting these processes result in a more extensive phenotypic disintegration in only one of the sexes.

A possible explanation for the lack of a role of host plant quality as a cue of seasonal progression, at least in thermal conditions of the dry season, could be that temperature suffices as a cue, as our lab population of *B. anynana* originated from a location in Malawi where temperature is a highly reliable predictor of seasonal transitions (Oostra et al. 2018). Thus, the reliability of temperature may override the necessity for additional cues in this population. Hence, an interesting open question is whether food quality may be a more important cue in other parts of the species’ range, where the relevance and reliability of temperature as a cue is less (Roskam and Brakefield 1996, van Bergen et al. 2017). Future studies could investigate the effect of host plant quality on life-history traits in populations or species originating from regions where temperature is a relatively unreliable cue. In addition, here we only exposed the larvae to old maize plants during the last two instars of development, while the larvae of this species utilize a variety of grass species in the wild and are likely to feed on poor quality host plants for longer periods (Brakefield et al. 2009, van Bergen et al. 2016). The deteriorating effects of the dry season conditions on natural grasses are likely to be host specific and longer exposure to poor quality host during development may trigger more pronounced phenotypic effects.

Taken together, plant quality affected life history traits in a temperature- and sex-specific manner in our study, indicating that under certain environmental condition a single cue (e.g. temperature) might suffice to shape an organisms’ phenotype, while under other conditions additional cues (like plant quality) might also become relevant in shaping the organism’s phenotype. Thus, it is important to study phenotypic plasticity in a multivariate environment as multiple environmental factors can interact to produce different phenotypes, and are better representative of conditions occurring in the wild.

## Acknowledgements

We would like to thank Andrew Balmer for help with the plant measurements, and the Radiating Butterflies Group for helpful discussions. This study was supported by funding for summer studentship from Trinity College, Cambridge, an INSPIRE scholarship travel fund to PS, and an Advanced Grant from the European Research Council (EMARES – 250325) to PMB.

